# *APOE4* poses opposite effects of plasma LDL on white matter integrity in older adults

**DOI:** 10.1101/2023.10.24.563796

**Authors:** Zhenyao Ye, Yezhi Pan, Rozalina G. McCoy, Chuan Bi, Mo Chen, Li Feng, Jiaao Yu, Tong Lu, Song Liu, Si Gao, Kathryn S. Hatch, Yizhou Ma, Chixiang Chen, Braxton D. Mitchell, Paul M. Thompson, L. Elliot Hong, Peter Kochunov, Tianzhou Ma, Shuo Chen

**Affiliations:** Maryland Psychiatric Research Center, Department of Psychiatry, School of Medicine, University of Maryland, Baltimore, Maryland, United States of America; Division of Biostatistics and Bioinformatics, Department of Epidemiology and Public Health, School of Medicine, University of Maryland, Baltimore, Maryland, United States of America; Division of Endocrinology, Diabetes, & Nutrition, Department of Medicine, School of Medicine, University of Maryland, Baltimore, Maryland, United States of America; Beth Israel Deaconess Medical Center, Harvard Medical School, Boston, Massachusetts, United States of America; Department of Nutrition and Food Science, College of Agriculture & Natural Resources, University of Maryland, College Park, Maryland, United States of America; Department of Mathematics, University of Maryland, College Park, Maryland, United States of America; School of Computer Science and Technology, Qilu University of Technology (Shandong Academy of Sciences), Jinan, Shandong, China; Department of Medicine, University of Maryland School of Medicine, Baltimore, Maryland, United States of America; Imaging Genetics Center, Keck School of Medicine, University of Southern California, Marina del Rey, California, United States of America; Department of Epidemiology and Biostatistics, School of Public Health, University of Maryland, College Park, Maryland, United States of America

**Keywords:** *APOE4*, cognition, metabolomics, moderation, white matter integrity

## Abstract

**INTRODUCTION:** APOE4 is a strong genetic risk factor of Alzheimer’s disease and is associated with changes in metabolism. However, the interactive relationship between APOE4 and plasma metabolites on the brain remains largely unknown.

**MEHODS:** In the UK Biobank, we investigated the moderation effects of APOE4 on the relationship between 249 plasma metabolites derived from nuclear magnetic resonance spectroscopy on whole-brain white matter integrity, measured by fractional anisotropy using diffusion magnetic resonance imaging.

**RESULTS:** The increase in the concentration of metabolites, mainly LDL and VLDL, is associated with a decrease in white matter integrity (b= -0.12, CI= [-0.14, -0.10]) among older *APOE4* carriers, whereas an increase (b= 0.05, CI= [0.04, 0.07]) among non-carriers, implying a significant moderation effect of *APOE4* (b= -0.18, CI= [-0.20,-0.15]).

**DISCUSSION:** The results suggest that lipid metabolism functions differently in *APOE4* carriers compared to non-carriers, which may inform the development of targeted interventions for *APOE4* carriers to mitigate cognitive decline.

## Introduction

Alzheimer’s disease (AD) is characterized by neuronal loss as well as the accumulation of beta-amyloid (Aβ) and tau in the brain and cerebrospinal fluid. The ε4 allele of the apolipoprotein E (*APOE*) gene stands as the most prominent genetic risk variant for late-onset AD, exerting a dose-dependent influence on disease susceptibility and progression (*APOE4/4* > *APOE3/4* > *APOE3/3*)^3^. *APOE* with isoform variations at amino acids position 112 (specifically, arginine for apoE4 and cysteine for apoE3) plays a critical role in lipid transport and the maintenance of cholesterol homeostasis via binding to metabolic receptors including the low-density lipoprotein receptor (LDLR) and lipoprotein receptor-related protein 1 (LRP1)^4^. Individuals with late-onset AD carrying the *APOE4* genotype have been reported to exhibit low levels of phosphatidylcholine (PC)^5^, and experience abnormal membrane functions, including impaired synaptic transmission and disrupted processing of the amyloid precursor protein, potentially contributing to the production of Aβ^6^.

AD involves complex neurophysiological changes, contributing to cognitive impairment, memory loss, disorientation, and behavioral changes^7^. Neuroimaging techniques have been commonly used to characterize AD, offering the capability to visualize and analyze detailed structural changes and neurodegeneration associated with AD in vivo^8^. White matter (WM) degeneration is regarded a pathological hallmark of various neurodegenerative diseases, particularly for AD^9^. WM integrity can be measured by fractional anisotropy (FA) using diffusion tensor imaging (DTI)^6^. Reductions in FA are indicative of neuronal fiber loss, degeneration of myelin and WM atrophy, making it a valuable marker in studying AD and its associated pathophysiology underlying AD^10^.

The impact of *APOE4* on WM changes, irrespective of AD, remains a subject of ongoing controversy, despite its established role in increasing the lifetime risk for AD in a dose-dependent manner. Williams et al.^11^ showed that cognitively normal adults carrying the *APOE4* genotype, as observed in the Baltimore Longitudinal Study of Aging, have a greater decline in WM integrity measured by FA in the genu and splenium of the corpus callosum, compared to non-carriers. Conversely, Lyall et al.^12^ found no association between the *APOE4* genotype and compromised WM tract integrity in older-aged adults examined within the UK Biobank (UKB) cohort.

Brain WM is composed of numerous bundles of axons that facilitate the transmission of electrical signals across various regions of the brain and spinal cord^13^. Axons are enveloped and insulated by a fatty substance known as myelin, which is predominantly composed of lipids, accounting for approximately 70-80% of its content^14^. *APOE4* has an impact on critical metabolic pathways, which can in turn influence WM. In a recent study, *APOE4* was associated with reductions in metabolites involved in the tricarboxylic acid (TCA) cycle, which is particularly important for predicting AD in the WM^18^. Additionally, the plasma metabolites profiles of *APOE4* showed altered neuronal sterol metabolism - characterized by reduced cholesterol and phospholipid secretion, decreased lipid-binding capacity, and increased intracellular degradation^19^.

Therefore, based on the evidence that metabolites and *APOE4* genotypes are both involved in the process of cognitive decline and deficits in WM integrity, this study aimed to examine the role of *APOE4* in the relationship between NMR-based plasma metabolic alterations and brain WM integrity. WM integrity was measured by FA in a large cohort of individuals aged 65 or older to gain insights into the potential to optimize dietary and lifestyle choices and thereby delay the onset of AD. By integrating genetic, neuroimaging, and metabolite data from 1,917 participants from the UKB, we tested a Gene-by-Metabolite (GxM) interaction hypothesis on how *APOE4* modifies the effect of plasma metabolites on regional WM tracts. We also investigated whether elevated levels of lipid metabolic biomarkers contribute to age-related cognitive decline, particularly in *APOE4* carriers.

## Materials and Methods

### Study sample and exclusion/inclusion criteria

The data analyzed in this study were sourced from the UKB (http://www.ukbiobank.ac.uk/)^20^, which obtained ethical approval from the National Information Governance Board for Health and Social Care and the National Health Service North West Multicenter Research Ethics Committee (REC reference 21/NW/0157). All UK Biobank participants provided written informed consent before participating in the study.

The UKB is a large prospective study that includes ∼500,000 participants aged 37-73 years at recruitment in 2006-2010 at 22 assessment centers across UK. At the initial assessment visit (2006-2010), the UKB collects medical history, lifestyle information, and biological samples. At the imaging follow-up visit (2014 and after), the UKB started to perform brain MRI assessments of a subset of ∼100,000 participants^21^.

In this study, our sample comprised of 1,917 individuals 1) aged 65 years or older at the imaging visit, 2) carrying *APOE3/3*, *APOE3/4* and *APOE4/4* combinations, and 3) having complete NMR-based plasma metabolomic data and WM FA outcomes, while excluding 1) non-Europeans, 2) related individuals, 3) those with brain injury, neurologic disorders, or mental illness based on ICD-10 (see more details in Supplementary Table 1), 4) with more than 2% missing genotypes and an *APOE* ε2 allele or 5) using medication for lowering cholesterol, or for diabetes, or taking exogenous hormones (i.e., oral contraceptive pill or minipill and hormone replacement therapy) based on UKB phenotypes 6177 and 6153. Figure 1 demonstrates the primary stages involved in sample selection. The baseline characteristics of the full UKB cohort and our study sample are summarized in Table 1.

**Figure 1.**
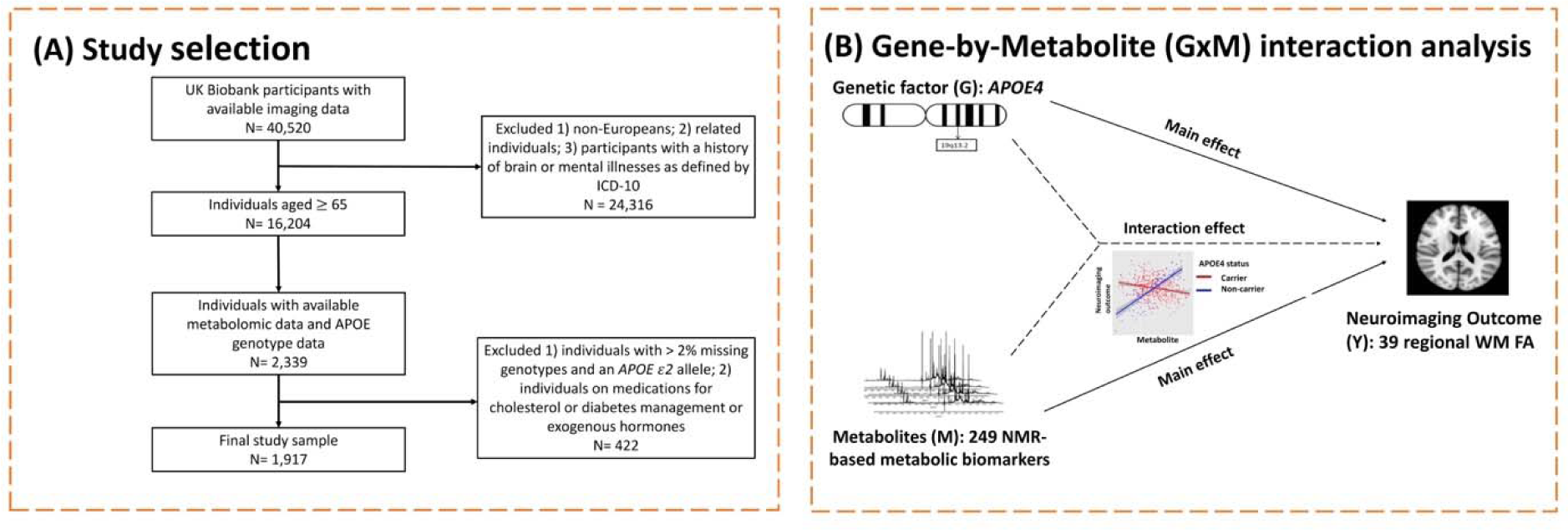
Study design. Panel (A) presents the flow of participants in the study. **Panel (B)** depicts the main analysis, focusing on the gene-by-Metabolite (GxM) interaction analysis. The *APOE4* genotype serves as the genetic factor (G), while 249 nuclear magnetic resonance (NMR)- based metabolic biomarkers are briefly called metabolites (M). The neuroimaging outcome (Y) is measured by fractional anisotropy (FA) in 39 regional white matter (WM) tracts. The main effects of G and M, as well as their interaction effect between G and M on Y, are examined through moderation analysis.

**Table 1.**
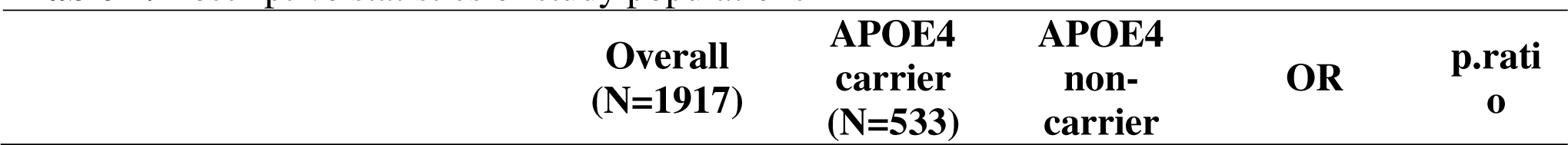
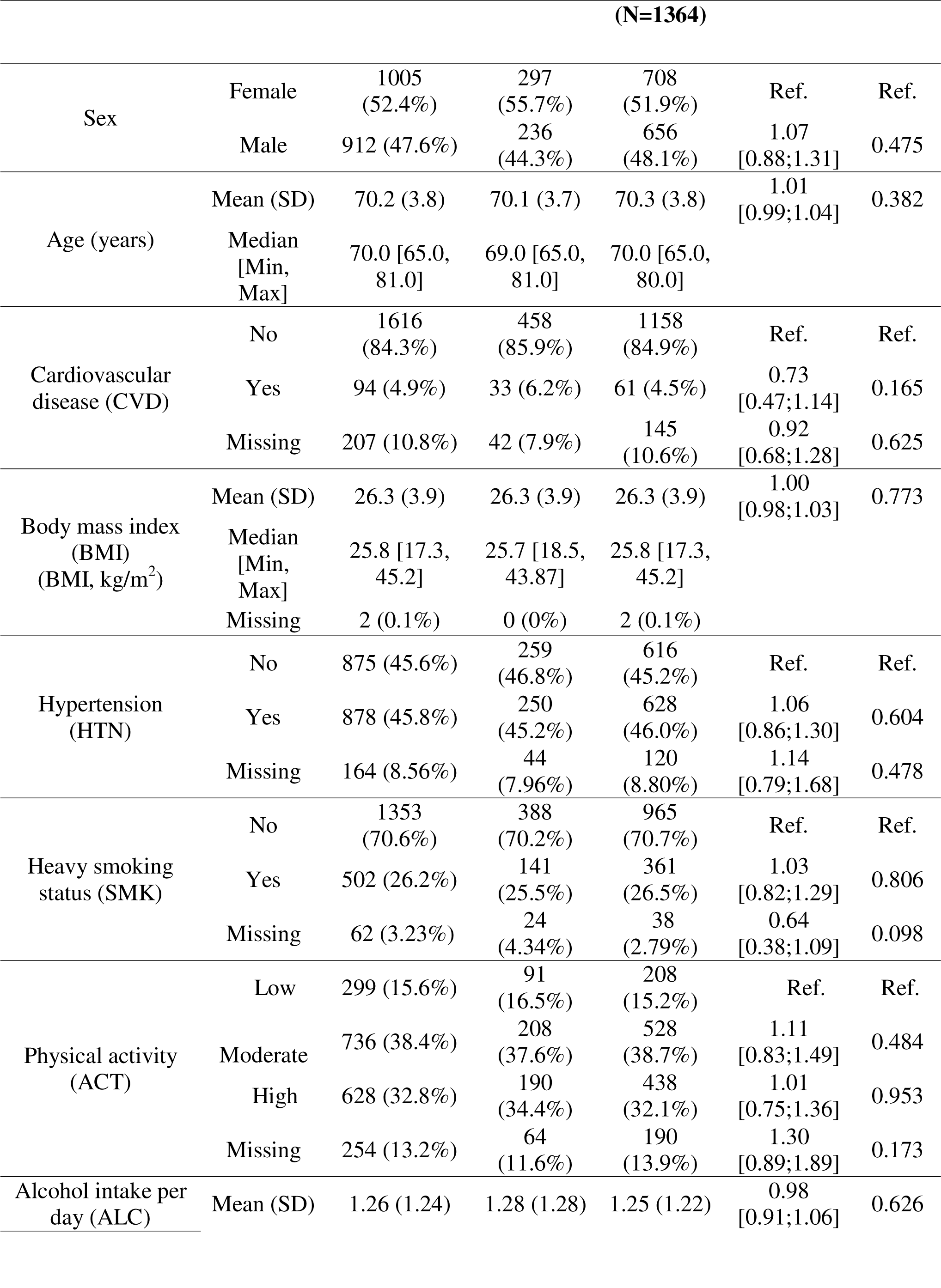
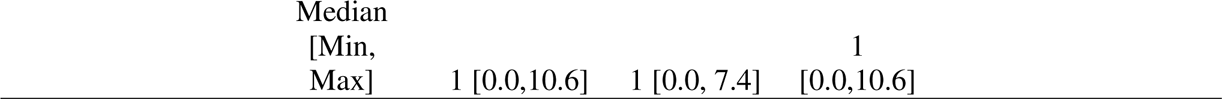
Descriptive statistics of study populations.

### *APOE* genotype data

Approximately 500,000 individuals from the UKB study were genotyped using two similar genotyping arrays, the Affymetrix UK BiLEVE Axiom and UK Biobank Axiom^TM^ platforms, to capture a comprehensive set of over 90 million single nucleotide variants (SNVs, detailed protocol available elsewhere^22^). Our analysis focused on unrelated individuals of European ancestry, predominantly of British and Irish descent. Furthermore, we excluded individuals with more than 2% missing genotypes using PLINK^23^ (version 1.9, www.cog-genomics.org/plink/1.9/). To determine the *APOE* genotype of participants, two *APOE* isoform coding single nucleotide polymorphisms (SNPs), rs429358 and rs7412, located on chromosome 19, were genotyped^24^. Individuals were categorized as *APOE4* carriers if they possessed the *APOE3/4* or *APOE4/4* combinations, while those with the *APOE3/3* were classified as non-carriers of the *APOE*4.

### Nuclear magnetic resonance-based metabolic biomarkers

In the UKB, Nightingale Health Plc measured metabolic biomarkers using high-throughput NMR spectroscopy in a randomly selected subset of non-fasting baseline plasma samples from 118,466 participants in the UKB cohort^25^. The final metabolic biomarker profiling underwent quality control procedures and encompassed a total of 249 metabolic measures, covering routine lipids, lipoprotein subclasses, fatty acid composition, as well as low-molecular-weight metabolic biomarkers (see full list of biomarkers in Supplementary Table 2). To ensure comparability and minimize biases, the metabolic biomarker values were normalized to the same range prior to analysis.

### White matter integrity data

WM integrity was calculated using diffusion tensor imaging (DTI) data processed by the UKB team, following the documented acquisition protocol^21^. To establish a standard-space white-matter skeleton, the DTI data underwent processing using the ENIGMA protocol. This involved averaging skeletonized images across a set of 48 standard-space tract masks and applying Tract-Based Spatial Statistics (TBSS) analysis. We focused on a total of 39 regional WM integrity using FA measure (see the complete list of WM FA measures in Supplementary Table 3), which represents a scalar value ranging from 0 to 1, providing insights into the extent of anisotropy within a diffusion process. A higher FA value indicates that the localized WM fiber bundles are more intact, suggesting a greater likelihood of water diffusion along the longitudinal axis of the WM bundles. To mitigate potential biases, individuals with brain illnesses, injuries, or mental disorders (as categorized by ICD-10, see more details in Supplementary Table 1) were excluded from the analysis.

### Cognitive g factor

Individuals underwent comprehensive cognitive assessments as a part of the UKB imaging study^26^. Our analysis focused on nine cognitive tests corresponding to seven specific domains: processing speed, perceptual reasoning, visuospatial learning/memory, cognitive flexibility, executive function/planning, working memory, and fluid intelligence. Prior to analysis, we conducted quality control on the cognitive data (see more details in Supplementary Table 4) and addressed missing data using multiple imputations with the predictive mean matching (PMM) method. We then derived a latent general intelligence factor (g factor) to provide a comprehensive measure of overall cognitive performance, capturing shared variance across these seven cognitive domains.

### Covariates

We included several covariates in our analyses to mitigate potential bias and account for confounding effects: age, age^2^, sex, body mass index (BMI), hypertension status (HTN, denoted as yes/no based on anti-hypertensive drugs use or systolic blood pressure >= 140 mmHg or diastolic blood pressure >= 90 mmHg), heavy smoking status (SMK, defined as individuals reporting smoking on most or all days in the past), cardiovascular disease (CVD, diagnosed using ICD-10 codes I60-I69, denoted as yes/no), average alcohol intake per day (ALC), and physical activity levels (ACT) based on the International Physical Activity Questionnaires (IPAQ). Missing data were identified in those covariates, exhibiting varying degrees of impact (CVD 207 [10.8%], BMI 2 [0.1%], HTN 164 [8.6%], SMK 62 [3.2%], ACT 254 [13.2%]). To address the potential bias introduced by missing values and enhance the accuracy of statistical estimates, we performed multiple imputations to impute missing data, and all subsequent analyses were performed on the complete datasets.

### Moderation analysis

We examined the main effects of *APOE4* and each metabolic biomarker on each WM tract FA measure, adjusting for the aforementioned covariates. Then, we investigated the GxM interaction effects between each metabolite and *APOE4* status (*APOE4* carriers vs. non-carriers) on each WM tract FA measure, by using the following linear regression model:

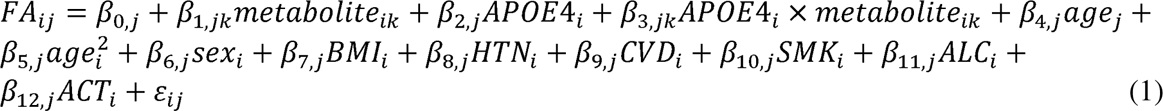

where *i* indicates individual, *i* = 1,2, …,1917; *j* indicates the specific WM FA tract, *j* = 1, 2, …, 39; *k* represents the specific metabolic biomarker, *k* = 1,2, …, 249; FA_*ij*_ represents the FA measure of the ith individual from the jth WM tract; metabolite_ik_ represents the value of the kth metabolic biomarker of the ith individual; the intercept *β*_*O,j*_, error term ε_*ij*_ and coefficients /3s represent the regression coefficients associated with each variable in the model. The *β*_3,jk_ is the main parameter of interest, corresponding to the GxM interaction effect in the model.

### Statistical analysis for multivariate metabolomics and brain imaging measures

Our moderation analysis involves multivariate outcomes (i.e., FA measures), multivariate metabolites, and a binary variable indicating the *APOE4* genotype status (*APOE4* carriers vs non-carriers). Multivariate-to-multivariate analysis is essential for required to detecting the underlying moderation patterns. Conventional canonical correlation analysis tools are not well-suited for our study because our focus is on identifying moderation patterns rather than multivariate-to-multivariate correlations. To address this need, we employ adaptive dense bipartite subgraph extraction algorithms^27^. The input for the algorithm is derived from the inference result matrix obtained through the above analysis, as defined by Eq (1). To identify the moderation pattern, our goal is to identify a dense maximal biclique consisting of a subset of metabolites and a subset of FA measures. Each metabolite along with the *APOE4* genotype status, which demonstrates a strong interaction effect on any FA measure, revealing moderation patterns (see Appendix). We further perform statistical inference for each detected biclique while controlling family-wise error rate by permutation tests.

Based on the results of multivariate analysis, we conducted a meta-analysis to examine GxM interaction effects between selected metabolic biomarkers and each chosen WM tract FA measure and demonstrate the meta-analyzed GxM interactions using forest plots.

### Age-related cognitive decline in different GxM subgroups

The selected metabolic biomarkers were further subjected to a factor analysis to generate a factor score. Additionally, a group variable was created to classify individuals into different levels of metabolites based on a median threshold of this factor score. Individuals with a factor score above the median were categorized as having a “high” level of metabolites, while those with factor scores below the median were classified as having a “low” level of metabolites. To explore the relationship between metabolic levels, *APOE4* status, and changes in cognitive performance over time, we compared the age trajectories of cognition in individuals with distinct metabolic levels and different *APOE4* statuses by using stratified analysis.

### Sensitivity analyses

To assess the impact of medication usage and ethnic heterogeneity on our primary findings, we conducted sensitivity analyses using the following criteria. First, we added additional individuals who self-identified with non-European ancestries (defined by UKB phenotype 21000) and who met our previous inclusion and exclusion criteria. Next, we included individuals who reported medication use for cholesterol, diabetes, or exogenous hormone intake based on UKB phenotypes 6177 and 6153. For each sensitivity analysis subset, we replicated the moderation analyses and employed the biclique approach.

All statistical models incorporated the same set of covariates. To address missing data, we used the R package ‘mice’ (v3.15.0)^28^ for multiple imputations. For each imputed dataset, we fitted the model of interest, and then combined the estimates from each model, along with their corresponding standard errors, using Rubin’s rules^29^, resulting in a single set of estimates that accounted for the imputation uncertainty. For factor analysis, R package ‘psych’ (v2.3.3)^30^ was employed.

## Results

### Participant characteristics

Among eligible participants, 533 individuals (297 [55.7%] women) were *APOE4* carriers and 1,364 individuals (708 [51.9%] women) were non-carriers of *APOE4*. The mean age of the *APOE4* carriers was 70.1 years (SD = 3.7 years), and of non-carriers was 70.3 years (SD = 3.8 years). There were no significant differences in the demographic characteristics of *APOE4* carriers and non-carriers (all *P* > 0.05; Table 1).

### The effects of *APOE4* on WM integrity

We first examined the main effect of *APOE4* on each regional WM FA tract (a total of 39 separate analyses). Each model was adjusted for age, age^2^, sex, BMI, CVD, HTN, SMK, ALC and ACT. Our study did not reveal any statistically significant main effects of *APOE4* on WM integrity for any of brain regions in our study (Benjamini-Hochberg adjusted P > 0.05, Supplementary Table 5).

### The effects of metabolites on WM integrity

We then investigated the main effect of each metabolite on each tract’s regional WM FA by pooling both *APOE4* carriers and non-carriers. A total of 249X39 analyses were performed, adjusting for the aforementioned covariates in the models. We did not observe any statistically significant results (Benjamini-Hochberg adjusted P> 0.05, Supplementary Table 6).

### Gene-by-Metabolite interaction effects on WM integrity

While these results suggested that there is no direct impact of either *APOE4* or metabolites on WM degeneration, it is possible that the interactions between *APOE4* and metabolites were not captured when analyzing the associations between *APOE4* and WM integrity, and between metabolites and WM integrity separately. To explore this further, we proposed a hypothesis suggesting the presence of GxM interactions, where the *APOE4* genotype modifies the effects of metabolites on brain WM integrity.

We performed moderation analyses to examine our hypothesis by regressing each regional WM FA measure on each plasma metabolites while considering its interaction with *APOE4* status (carriers vs non-carriers; see methods section for details). These analyses resulted in a total of 249×39 *p*-values, representing the interaction term for each pair. We then transformed the *p*-values into their negative logarithmic form (-log (p)) and employed a biclique approach (for details see methods section) to identify dense and cohesive patterns between metabolites and FA (permutation test *p*-value < 0.001).

We found 25 metabolic biomarkers with strong GxM interaction effects on six WM FA tracts (see Figure 2), including the bilateral anterior limb of the internal capsule (ALIC-L/-R), bilateral posterior limb of the internal capsule (PLIC-L/-R), splenium of the corpus callosum (SCC), and the right hemisphere of the superior longitudinal fasciculus (SLF-R). Among these regions, SCC carries fibers that accept visual stimuli and facilitate auditory processing^31^. PLIC integrates sensory information and the execution of motor functions^32^; ALIC facilitates decision-making and integration of signals related to emotion, motivation and cognition^33^; SLF contributes to specific functions in motor regulations, visuospatial attention, and somatosensory information processing within the brain . The 25 metabolic biomarkers in the detected submatrix mainly belong to the LDL (*n*=20) and VLDL (*n*=3) classes (see Figure 2). Figure 3 presents the effect sizes (symbolized by circles) of these 25 metabolites on the six WM tracts among *APOE4* carriers (*β_1,jk+_β_3,jk_*), among *APOE4* non-carriers (*β_1,jk_*) and the interaction effect (*β_3,jk_*), respectively (see details for /3 in Equation 1, methods section). To ensure fair comparisons, all effect sizes were standardized by dividing them by the standard deviation (SD) of each selected WM FA measurement. We found that all these metabolites have consistently negative effects on WM FA among *APOE4* carriers (*yellow* color) but positive effects among *APOE4* non-carriers (*purple* color), resulting in negative interaction effects (i.e., difference in main effect; *blue* color). The meta-analyzed results over all metabolites (symbolized by diamonds) are significant and consistent with individual results (*P* < 0.0001 for all three types of effect sizes). Specifically, the meta-GxM interaction is associated with a 20.2% decrease in FA within ALIC-L (CI = 17.9, 22.5), 18.1% within ALIC-R (CI = 15.8, 20.5), 15.8% within PLIC-L (CI =13.4, 18.2), 17.8% within PLIC-R (CI =15.3, 20.2), and

**Figure 2.**
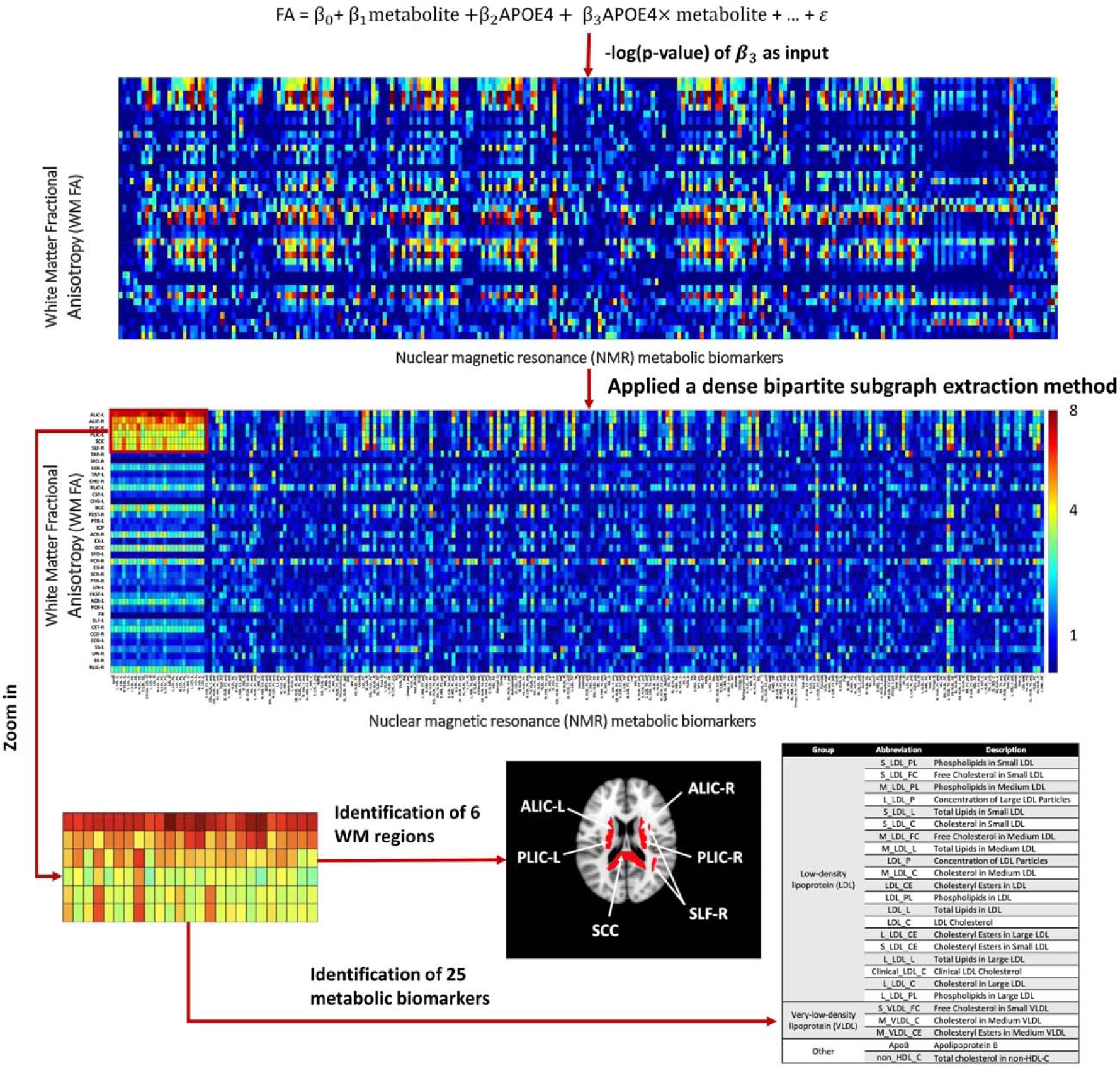
A representation of dense and cohesive metabolite by FA patterns. A total of six white matter (WM) tracts and twenty-five metabolic biomarkers were identified through a dense bipartite subgraph extraction method utilizing the negative logarithm of p-values of the coefficients /3_3_ of the interaction term (*APOE4*Xmetabolite, see Equation 1 in the methods section, ‘Moderation analysis’, for details). The six identified WM tracts include the bilateral anterior limb of internal capsule (ALIC-L/-R), bilateral posterior limb of internal capsule (PLIC- L/-R), splenium of corpus callosum (SCC) and right hemisphere of superior longitudinal fasciculus (SLF-R). The twenty-five metabolites shown in the table were mainly categorized into groups low-density lipoprotein (LDL, *n* = 20) and very-low-density lipoprotein (VLDL, *n* = 3). Additionally, apolipoprotein B and total cholesterol in non-HDL-C were classified into another group.

**Figure 3.**
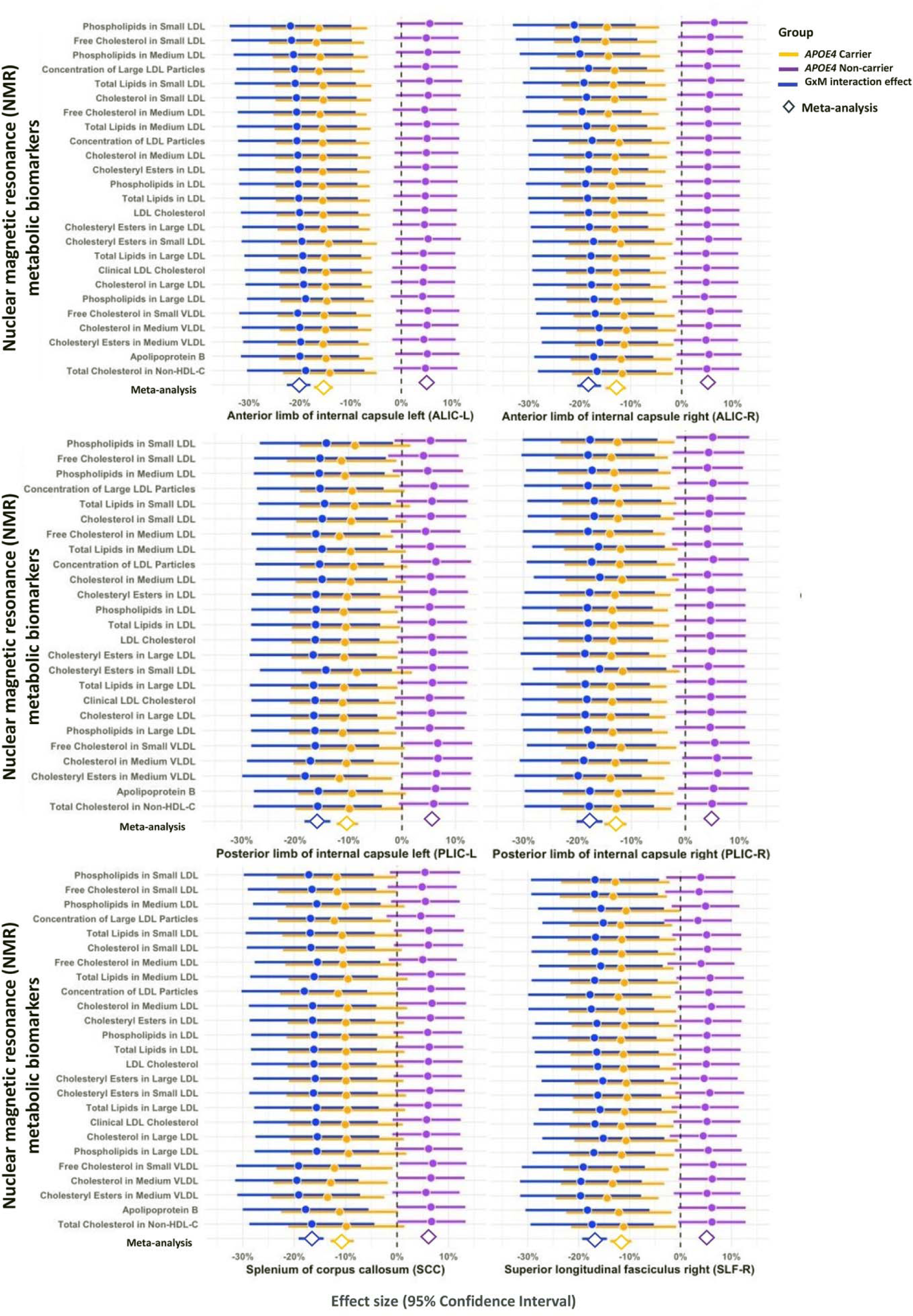
Forest plots of metabolite effect sizes on white matter (WM) tracts. The effect sizes are represented by circles, indicating the magnitude of effect for each metabolite on each WM among *APOE4* carriers (*β_1,jk_+β_3,jk_*), depicted in *yellow*), among *APOE4* non-carriers (*β_1,jk_)* depicted in *purple*) and interaction effect *β_3,jk_*, representing the difference in main effect, depicted in *blue*) (see Equation 1 in the Methods section for details). We standardized the effect sizes by dividing them by the standard deviation (SD) of each identified WM FA measure for a consistent assessment of the magnitude of the effects across the different WM tracts. We calculated meta-analyzed effects (*P* < 0.001) of all metabolites on WM FA, which are illustrated as diamonds, providing an overall estimate of the collective impact of metabolites on WM integrity.

16.7% within SCC (CI = 14.3, 19.1) and 16.9% within SLF-R (CI = 14.5,19.3) (see Figure 3). The cross region meta-analysis results indicate that an increase in the concentration of metabolites is associated with a 12% decrease in WM (CI=[-0.14, -0.10]) among older APOE4 carriers, while a 5% increase (CI=[0.04, 0.07]) among non-carriers. This implies a significantly negative moderation effect of APOE4 (/3 = -0.18, CI=[-0.20,-0.15]).

### Sensitivity analyses

We performed several sensitivity analyses to show the robustness of our results. To mitigate the selection bias towards European participants, we pooled a sample of 1,954 individuals including both European and non-European ancestries to assess the influence of ethnic heterogeneity on our primary results. The identified brain regions and metabolites as illustrated in Supplementary Fig. 1 (A) in Appendix, showed minimal changes, suggesting a consistent pattern using (-log(p)) values of GxM interaction terms. Furthermore, we broadened our analysis to include the 422 individuals who had reported medication usage for cholesterol, diabetes, or exogenous hormones, resulting in a larger cohort totaling 2339 individuals. The outcomes of this sensitivity analysis, depicted in Supplementary Fig. 1 (B) in Appendix, also revealed a dense FA- metabolomics pattern that consistently aligns with our primary findings.

### Age-related cognitive decline in different GxM subgroups

We further delved deeper into understanding how our findings are related to cognitive decline. We derived a latent variable that accounts for 70% of the total variance from the 25 identified metabolites, representing the overall level of metabolites and their collective impact. Participants were categorized into four groups based on their APOE4 status and metabolic levels, including 1) Group “Carrier-High”: 303 *APOE4* carriers with high metabolic levels; 2) Group “Carrier- Low”: 211 *APOE4* carriers with low metabolic levels; 3) Group “Non-carrier-High”: 589 APOE4 non-carriers with high metabolic levels; 4) Group “Non-carrier-Low”: 684 APOE4 non-carriers with low metabolic levels.

We observed that individuals who are *APOE4* carriers and have high levels of LDL/VLDL experience the most rapid cognitive decline with increasing age as compared to individuals in other groups (P = 0.008, see Figure 4). This finding confirms that the combination of carrying the *APOE4* and metabolic dysfunction with elevated levels of LDL/VLDL contributes to an accelerated cognitive decline over time.

**Figure 4.**
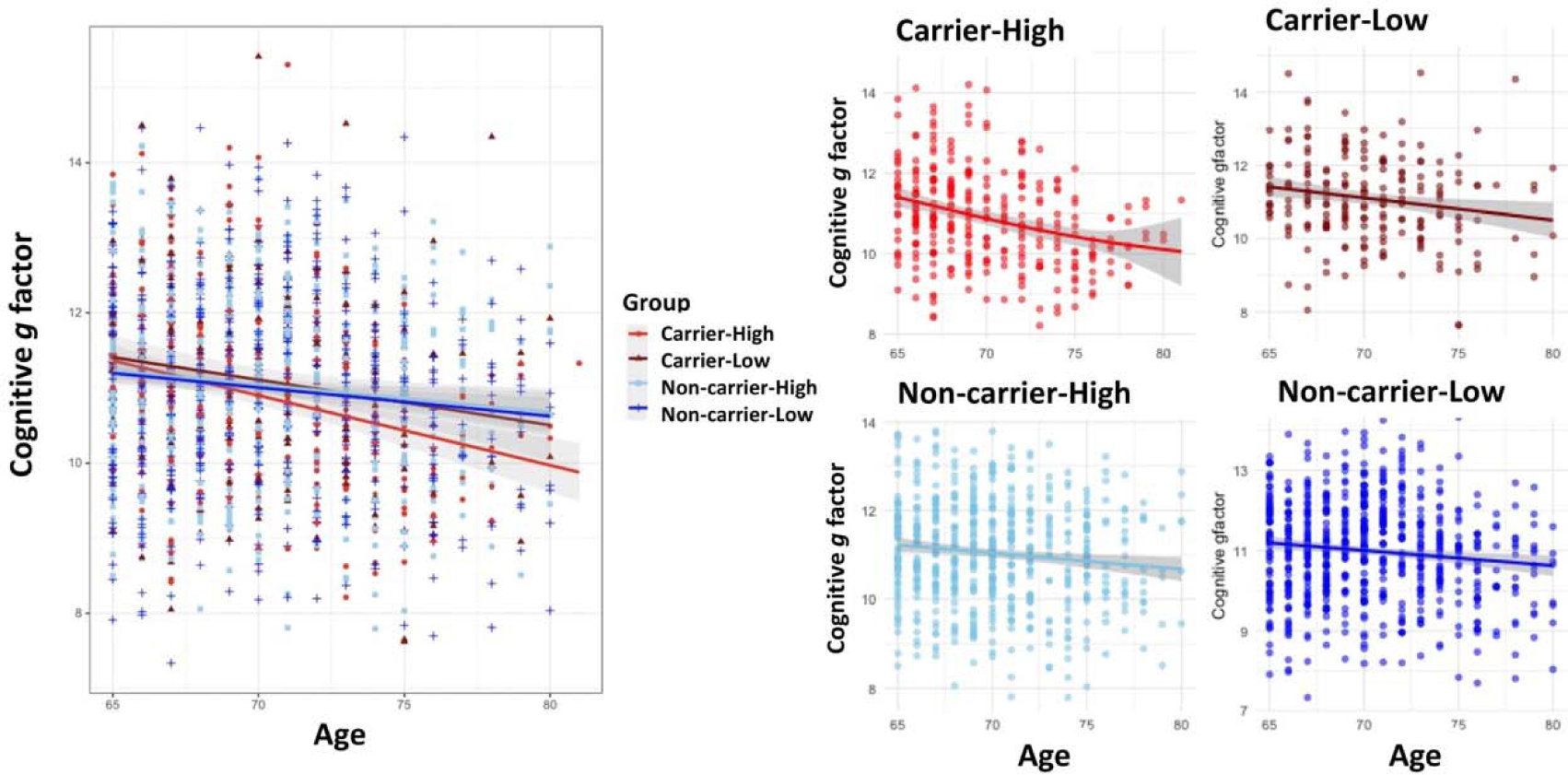
Impact of Gene-by-Metabolite (GxM) interaction effects on cognitive performance over age. Based on the combination of *APOE4* status (carriers vs. non-carriers) and metabolic levels (at a median threshold), we categorized study participants into four groups: “Carrier-High” (*red* color) represents *APOE4* carriers with high metabolic levels (above the median threshold); “Carrier-Low” (*brown* color) represents *APOE4* carriers with low metabolic levels (below the median threshold); “Non-carrier-High” (*skyblue* color) represents non-carriers with high metabolic levels and “Non-carrier-Low” (*blue* color) represents non-carriers with low metabolic levels.

## Discussion

In this study, we investigated the interaction effects of *APOE4* genotype and plasma metabolites on regional WM integrity by integrating genetic, neuroimaging and NMR data from a large UKB cohort. While we did not find a direct main effect of *APOE4* and metabolites on WM degeneration, our results revealed a significant interaction between *APOE4* and metabolites on WM integrity. Specifically, we found elevated levels of LDL/VLDL have a detrimental impact on WM integrity among *APOE4* carriers but not among non-carriers. These findings provide valuable insights into the underlying biological mechanisms involved in the development of AD, help identify individuals at a higher risk, enhance the accuracy of AD prediction and suggest potential interventions – lifestyle and pharmacologic – to prevent AD onset and progression.

We identified a set of 25 metabolic biomarkers whose effects on WM integrity are modified by *APOE4*. These metabolites are found in VLDL with diameters ranging from 36.8 nm to 44.5 nm as well as LDL with diameters ranging from 18.7 nm to 25.5 nm (Figure 2 and Supplementary Table 2). This is consistent with earlier studies highlighting VLDL as a triglyceride-rich lipoprotein that exhibits a strong affinity for binding with *APOE4*^30^. In the presence of peripheral (hepatocyte-derived) *APOE4*, there is a competition for binding between VLDL and LDL particles, which can affect the clearance and metabolism of both lipoproteins, consequently affecting their respective levels in the bloodstream^31^. This competition has been further supported by Garcia et al., who discovered that individuals who are very lean and carry the *APOE4* exhibit notably elevated levels of LDL compared to those with only the APOE3^32^. These findings suggest that peripheral *APOE4* enhances the competitive binding of VLDL to the LDL receptor, resulting in the subsequent elevation of plasma LDL levels.

Furthermore, in both peripheral and central (primarily astrocyte-derived) nervous systems, ApoE plays a crucial role in maintaining the integrity of the blood-brain barrier (BBB)^33^. *APOE4* genotype is associated with a compromised BBB, leading to increased permeability. This defective BBB allows a significant flux of cholesterol into the brain and/or loss of cholesterol from the brain into circulation^34^. LDL has been observed to be transcytosed through the BBB via a receptor-mediated mechanism, using specialized endocytic vesicles that avoid fusion with lysosomes^35^. High levels of LDL may increase the amount of oxidized LDLs^36^ that are able to pass the BBB into the brain, thereby potentially affecting neurological processes. The presence of high LDL and its oxidative modifications in the brain contribute to myelin loss and axon degeneration through various mechanisms, including the generation of oxidative stress and the initiation of an inflammatory response^37^. These processes can lead to abnormalities in white matter microstructure^11^ and have implications for decision-making and processing speed^38^.

VLDL and LDL are highly atherogenic due to their small size, high cholesterol content, and pro-inflammatory properties^39^. They penetrate the vascular wall and become trapped in the intima, leading to the accumulation of foam cells and the initiation of low-grade inflammation^40^, ultimately contributing to the development of atherosclerosis, cardiovascular, and cerebrovascular disease^41^. This diminished blood flow to the brain may hinder the supply of oxygen and nutrients to cerebral tissues, potentially contributing to cognitive impairment and the progression of AD^42^.

Our findings suggest that *APOE4* carriers with elevated LDL levels are at a higher risk of experiencing the transcytosis of LDL across the BBB, which can contribute to the development of atherosclerosis and synergistically impact cognitive decline. This process may also have implications for the integrity of myelin and axons, leading to degeneration of the WM. In contrast, individuals who do not carry the *APOE4* allele generally possess a healthier BBB that functions as a protective barrier, preventing the entry of peripheral LDL into the brain. Additionally, non-carriers exhibit reduced affinity of binding between VLDL and LDL with the LDLR resulting in a lower exchange of LDL between peripheral and cerebral areas. This limited exchange helps to maintain lipid homeostasis in the brain, further protecting against the potentially detrimental effects of elevated LDL. As depicted in Figure 4, our findings demonstrate that *APOE4* carriers with high LDL levels display heightened susceptibility to cognitive decline compared to non-carriers, further supporting our hypothesis and emphasizing that the significance of *APOE4* in modifying the effects of LDL/VLDL-related metabolites on WM integrity. However, future studies should incorporate more advanced methods and experimental designs to validate and build on these findings in biological settings. This should facilitate a deeper understanding of the underlying mechanisms involved and may uncover potential therapeutic interventions.

Metabolic dysfunction with obesity, hyperlipidemia, and hypertension are known risk factors for AD onset and progression^43^. While lipid-lowering therapy with statins has been associated with reduced risk of cognitive impairment and AD onset in observational studies^44^, this was not observed in randomized controlled trials, raising concerns that the association seen in observational studies is driven by confounding. Our study offers another possibility; specifically, that the association between dyslipidemia and cognitive impairment/AD is observed primarily in susceptible individuals with the *APOE4* genotype. This heterogeneity of treatment effects will need to be examined. Our study further provides a plausible link between vascular and degenerative cognitive decline in susceptible individuals.

To summarize, our study is a novel investigation of the GxM interaction between *APOE4* and metabolites on WM integrity. Our findings reveal that *APOE4* amplifies the adverse effects of metabolites associated with LDL/VLDL on specific WM tracts linked to motor, sensory, and executive functions. This amplification occurs by increasing the permeability of the BBB, allowing these metabolites to enter the brain. As a result, disturbances in lipid homeostasis, proinflammatory responses, and axonal loss are triggered, contributing to impairments in WM and processes related to myelin. The findings emphasize the importance of considering GxM interactions in disease prognosis and shed light on the metabolic changes that occur during the disease progression, particularly under the influence of specific genotypes.

## Acknowledgements

Data used in this study were obtained from the UK Biobank (https://www.ukbiobank.ac.uk/). We thank the UK Biobank cohort Group that collected the data made available for this work.

This study is supported by the National Institute on Drug Abuse of the National Institutes of Health.

## Funding

This research was funded by the National Institute on Drug Abuse of the National Institutes of Health under Award Number 1DP1DA04896801. Additional support for computer cluster was provided by NIH R01 grants EB008432 and EB008281.

## Competing interests

The authors report no competing interests.

## Notes

### Competing Interest Statement

The authors have declared no competing interest.

