## Appendix for "*APOE4* poses opposite effects of plasma LDL on white matter integrity in older adults"

**Supplementary Methods**

### Statistical analysis for multivariate metabolomics and brain imaging measures

The input for the algorithm is derived from the inference result matrix obtained through the moderation analysis, as defined by Eq (1) in the Method section. It forms in a $K\times J$ matrix denoted as W, where $K$ represents metabolites and $J$ denotes outcomes (i.e., FA measures). In this matrix, each entry $w_{kj}$ corresponds to values of $-$ $log(p_{kj})$, with $p_{kj}$ representing the p-values of the moderation effect of $k$th metabolite and *APOE4* genotype status on the $j$th FA measure. The matrix $W$ can be seen as the weight matrix of a weighted bi-partite graph $G[$ $U,$ $V, W]$ with two sets of nodes and weighted edges, where $U$ represents the first node set of metabolites and $V$comprises the second node set of the FA measures, and $W$ are weighted edges reflecting moderation levels. In cases where the moderation effect is more significant, the edge weight $w_{kj}$is higher.

Under the null hypothesis of no moderation effects, the high-weight edges are false positive findings and randomly distributed in $G[$ $U,$ $V, W]$. Recent advances in graph statistical theory have shown that a bi-partite subgraph, characterized by both denseness and considerable size, is less likely to arise from a null hypothesis of graph^1–3^. Identifying a dense and substantial bi-partite subgraph implies the validity of the alternative hypothesis, signifying the existence of multivariate-to-multivariate moderation. Therefore, to identify the systematic moderation pattern, our goal is to identify a dense maximal biclique consisting of a subset of metabolites, denoted as $S$ ($S\subseteq U$), and a subset of FA measures $T$ ($T\subseteq V$ $)$. Each metabolite in $S$ along with the *APOE4* genotype status, which demonstrates a strong interaction effect on any FA measure in $T$, revealing systematic moderation patterns. Computationally, we aim to detect the bicliques through optimizing the objective function $j_{\lambda}\left( S,T \right)$.

$\underset{S,T}{\mathrm{argmax}} j_{\lambda}\left( S,T \right)=\underset{S,T}{\mathrm{argmax}} \sum_{k,j} \sum_{k\in S, j\in T} [log(|w_{kj}|)-\frac{1}{2}\lambda log(|S||T|)$	(2)

where λ>1 is the tuning parameter that can be objectively determined by the Kullback–Leibler (KL) divergence criterion^4^.

**Supplementary Figures**

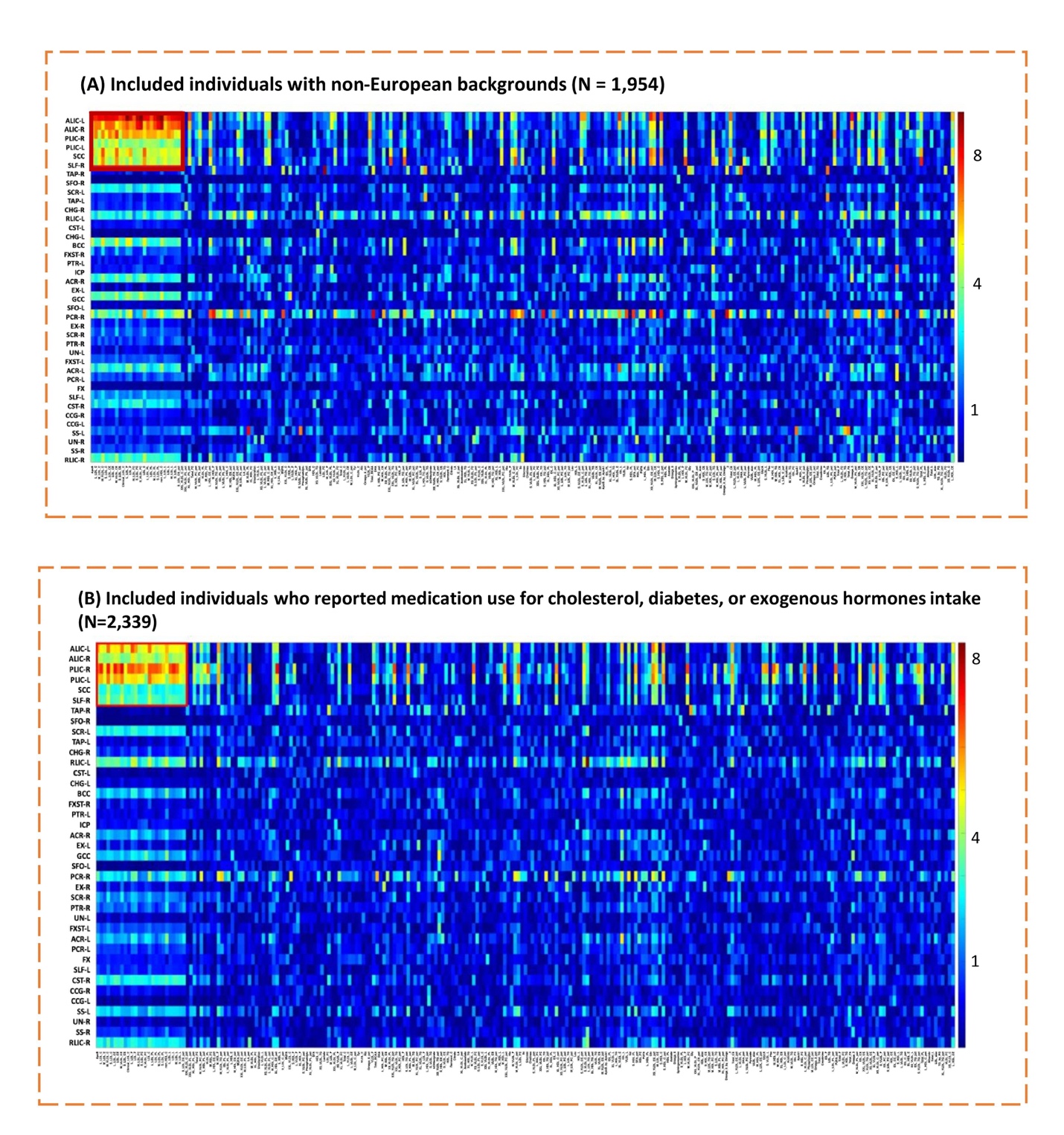

**Supplementary Fig. 1. Sensitivity analysis results.** A total of six white matter (WM) tracts and twenty-five metabolic biomarkers were identified through a dense bipartite subgraph extraction method utilizing the negative logarithm of p-values of the coefficients $\hat{\beta}_{3,jk}$ of the interaction term (APOE4$\times$metabolite, see Equation 1 in the methods section for details).
